# Environment Shapes the Accessible Daptomycin Resistance Mechanisms in *Enterococcus faecium*

**DOI:** 10.1101/607440

**Authors:** Amy G. Prater, Heer Mehtaa, Abigael J. Kosgei, William R. Miller, Truc T. Tran, Cesar A. Arias, Yousif Shamoo

## Abstract

Daptomycin binds to bacterial cell membranes and disrupts essential cell envelope processes leading to cell death. Bacteria respond to daptomycin by altering their cell envelopes to either decrease antibiotic binding to the membrane or by diverting binding away from vulnerable septal targets to remodeled anionic phospholipid membrane patches. In *Enterococcus faecalis*, daptomycin resistance is typically coordinated by the three-component cell-envelope-stress-response system, LiaFSR. Here, studying a clinical strain of multidrug-resistant *Enterococcus faecium* containing alleles associated with activation of the LiaFSR signaling pathway, we found that specific environments selected for different evolutionary trajectories leading to high-level daptomycin resistance. Planktonic environments favored pathways that increased cell surface charge via *yvcRS* upregulation of *dltABCD* and *mprF*, causing a reduction in daptomycin binding. Alternatively, environments favoring complex structured communities, including biofilms, evolved both diversion and repulsion strategies via *divIVA* and *oatA* mutations, respectively. Both environments subsequently converged on cardiolipin synthase (*cls*) mutations, suggesting the importance of membrane modification across strategies. Our findings indicate that *E. faecium* can evolve diverse evolutionary trajectories to daptomycin resistance that are shaped by the environment to produce a combination of resistance strategies. The accessibility of multiple and different biochemical pathways simultaneously suggests that the outcome of daptomycin exposure results in a polymorphic population of resistant phenotypes making *E. faecium* a recalcitrant pathogen.

## Introduction

The rise of multidrug resistant (MDR) pathogens is one of the most pressing biomedical problems of this century. The Center for Disease Control (CDC) reports that 2 million antibiotic resistant infections resulting in 23,000 deaths occur each year (1). Vancomycin-resistant enterococci (VRE) cause approximately 1,300 deaths annually with the number of infections increasing substantially over the last 15 years (1). The Infectious Disease Society of America has listed *E. faecium* (*Efm*) among the no ‘ESKAPE’ pathogens (***E****nterococcus faecium*, ***S****taphylococcus aureus*, ***K****lebsiella pneumoniae*, ***A****cinetobacter baumannii*, ***P****seudomonas aeruginosa*, ***E****nterobacter* spp.) for which there is an urgent need for new therapies (2). *Efm* accounts for a significant amount of enterococcal health-care-associated infections, particularly in severely immunocompromised patients. *Efm* strains that are resistant to all anti-enterococcal antibiotics have been widely described (3–6), making these infections untreatable in certain scenarios (1, 3–5, 7, 8).

DAP is a bactericidal cyclic lipopeptide antibiotic approved in 2003 and used widely as a “rescue” drug against MDR Gram-positive organisms such as *Staphylococcus aureus, Efm* and *Enterococcus faecalis* (*Efc*) (4, 9, 10). While the DAP mechanism-of-action remains unclear, DAP acts in a calcium-dependent-manner, where the DAP:Ca^+2^ complex binds to the cell membrane (CM) in a phosphatidylglycerol-dependent manner with high avidity for division septa. DAP binding has pleiotropic effects in the CM that include mislocalization of key proteins involved in cell wall and CM metabolism, and ultimately cell death (11–14). Unfortunately, DAP resistance is increasingly observed in the clinic. A clear understanding of the biochemical basis for resistance can potentially identify novel targets, therapeutic approaches, and diagnostic markers for these MDR infections to restore the effectiveness of antibiotics (including DAP) against recalcitrant strains (15).

To date, the manner in which organisms evolve DAP resistance falls into three categories: “repulsion” from the cell surface (16–18), “diversion” of DAP binding away from the division septa (13, 15, 16, 18–20) and antibiotic hyperaccumulation in selected cells (seen in streptococci) (21) to protect the bacterial population. *S. aureus* predominately exploits repulsion, often mediated through gain-of-function mutations in the *dltABCD* operon and the multiple peptide resistance factor, *mprF* (16, 18, 20). Mutations in the *dltABCD* operon, responsible for incorporating D-alanine into lipoteichoic acids (LTA), often result in a more positively charged cell envelope and reduced DAP binding (18, 22, 23). Similarly, MprF catalyzes the transfer of a lysyl group to phosphatidylglycerol causing a net increase in cell surface charge and reduced DAP binding (18). Although, it has also been postulated that this reaction affects DAP susceptibility by decreasing the availability of phosphatidylglycerol for DAP binding (24). In contrast, *Efc* employs a different strategy that involves redistribution of anionic phospholipid microdomains away from the septum, to divert DAP from critical septal targets; this mechanism is mediated by the LiaFSR cell-envelope-stress-response system in association with cardiolipin synthase (*cls*) (15, 25, 26). Our initial mechanistic studies (27) suggest that DAP resistance in *Efm* is not mediated by membrane remodeling but rather involves repulsion of the antibiotic from the cell surface. Nonetheless, the LiaFSR system is involved in the *Efm* DAP response since deletion of the gene encoding the LiaR response regulator resulted in hypersusceptibility independent of the genetic background of the strain (28).

To deconstruct the potential resistance strategies available to *Efm*, we used specific adaptive environments to evolve the clinical strain, *Efm* HOU503 that harbors the most common clinical LiaRS substitutions (LiaR^W73C^ and LiaS^T120A^), which predispose system activation, to DAP resistance (29). Despite HOU503 being poised to exploit potential redistribution pathways via LiaFSR activation, we show that we can use the environment to favor different biochemical strategies. Unlike the evolution of *Efc* and *S. aureus* to DAP resistance, our work suggests that *Efm* is able to employ varying and different resistance strategies that may help explain the recalcitrant nature of these infections and the therapeutic challenges they pose (27).

## Results

### Distinct adaptive environments select for divergent phenotypes and evolutionary trajectories

To favor distinct evolutionary trajectories that might be associated with the evolution of DAP resistance by *Efm* in the presence of LiaFSR substitutions, we performed experimental evolution using two different techniques that established a very different basis for the selection of adaptive phenotypes. We chose *Efm* HOU503 as the ancestor because it is a VAN resistant isolate (minimum inhibitory concentration (MIC) > 256 µg/ml) with a DAP MIC of 3 µg/ml and is poised to make the transition to clinical resistance (8). HOU503 contains mutated alleles within the LiaFSR pathway (LiaR^W73C^ and LiaS^T120A^) that in combination increase the strength of LiaFSR signaling (29). Thus, this strain provided the ideal scenario to investigate subsequent and viable evolutionary trajectories upon antibiotic exposures when an initial step has been taken.

First, we evolved five HOU503 populations to DAP resistance using a traditional serial-flask transfer model where planktonic cells were transferred daily to increasing DAP concentrations. The stepwise increase in DAP concentration was performed below the current population MIC to allow the establishment of multiple evolutionary trajectories within the population (25). The populations were passaged for a total of eight days with the final populations containing 8 µg/ml DAP (resistant by clinical standards). The environment established by flask-transfer strongly reduces the selection for trajectories that might rely upon biofilm as cells that adhere to surfaces are less likely to be transferred.

Next, we evolved two, independent HOU503 populations to DAP resistance in a bioreactor where the vessel remained constant and the culture was maintained at its fastest growth rate. As in the flask-transfer experiments, the bioreactor population was subjected to stepwise increases in DAP concentration, below the population MIC to maintain diversity, until the final population was growing at 8 µg/ml DAP, within 10 to 12 days. The bioreactor environment is, in many respects, the opposite of the flask-transfer environment, where cells that form biofilms or adhere to surfaces remain within the vessel, while planktonic cells, though still viable, are disadvantaged and can be washed out with a higher frequency. This production of biofilms contributes, in part, to the high level of polymorphism found within the bioreactor (Supplementary Text).

Following adaptation, two isolates from each of the five flask-transfer populations (10 isolates total) were selected for whole genome sequencing (WGS) and phenotypic characterization. From each bioreactor-adapted population, 10 and 9 isolates, respectively, with distinct phenotypic properties (DAP MIC, cell density at stationary phase, and the ability to form floc) underwent WGS and further characterization (19 isolates total). Antibiotic cross-sensitivities were also tested and are discussed in the Supplementary Text and Table S1.

To confirm that each technique produced a different adaptive environment, we performed a crystal violet assay to quantify end-point isolate biofilm growth and assayed the growth rates of 16 isolates with diverse genomes from both environments. As shown in Fig. 1A, bioreactor-derived isolates produced up to 10-fold more biofilm than HOU503 and the flask-transfer isolates, consistent with growing in different adaptive environments. While the bioreactor-derived isolates typically formed strong biofilms, isolate R2P29 produced little biofilm, consistent with our previous studies showing that bioreactors can favor highly polymorphic populations and maintain planktonic sub-populations (25, 30–33). This isolate is discussed further in the Supplementary Text.

**Fig. 1:**
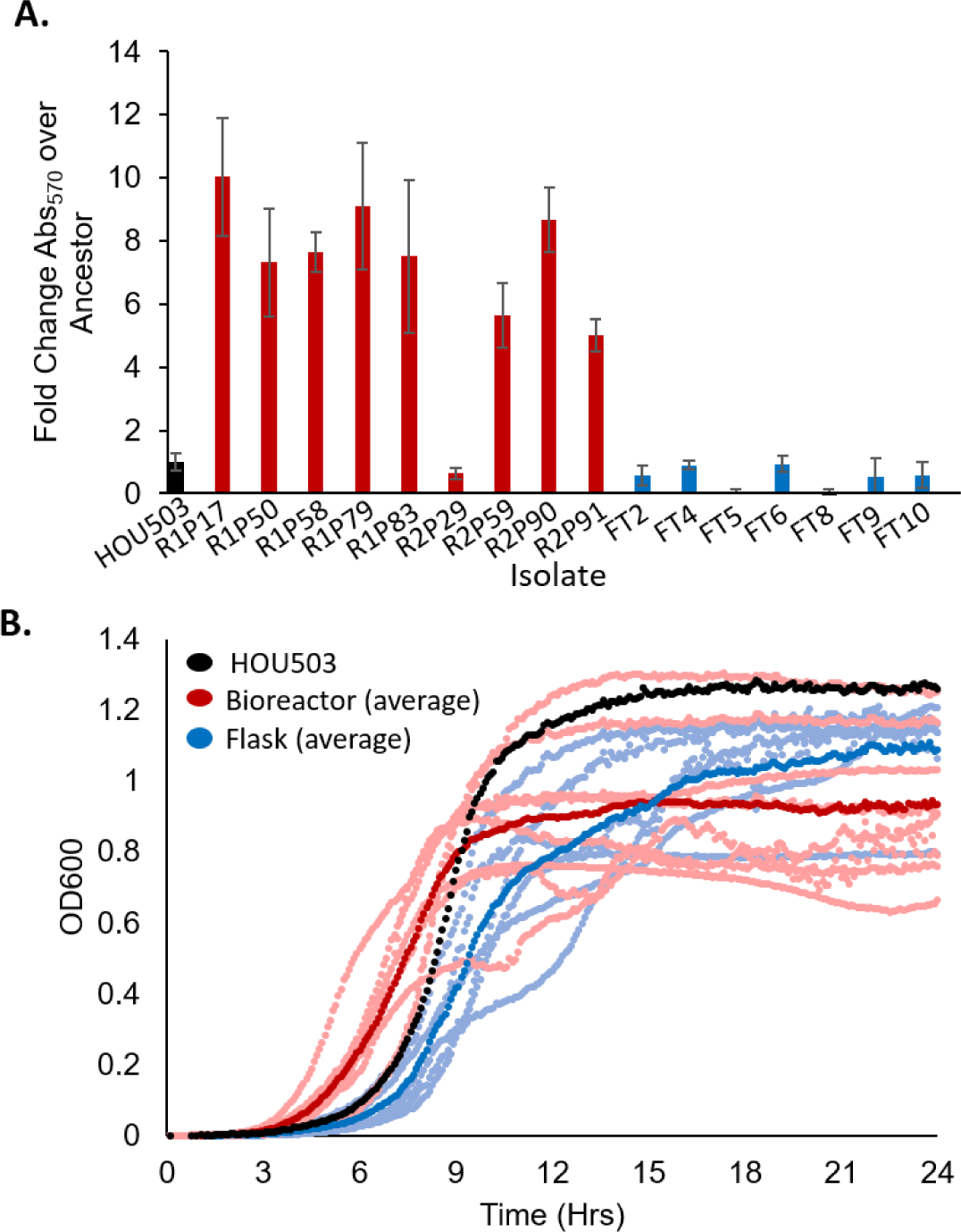
Varying the adaptive environment selects for distinctive and divergent phenotypes. The bioreactor (red) and flask (blue) environments evolve distinctly different phenotypes. **A.** A crystal violet assay was used to quantify biofilm formation of 16 end-point isolates and is reported as the fold change crystal violet Abs_570_ over the ancestor. All bioreactor isolates (except R2P29) produced significantly more biofilm than the ancestor (p<0.05) using two-sided T-test. Error bars represent standard deviation. **B.** Growth rates were performed in a microplate reader in triplicate for all end-point isolates. The dark red and blue markers indicate the average growth for each adaptive environment with the lighter shades showing the average growth of individual isolates.

In addition to biofilm formation, the growth rates of each end-point isolate type differed (Fig. 1B). Flask-transfer isolates grew more slowly than both the ancestor and the bioreactor-adapted isolates, taking 8.6-11.9 hours to reach the mid-point of their final cell density whereas HOU503 took 8.5 hours and the bioreactor-derived isolates took 5.5-8.9 hours (Fig. 1B). Since the bioreactor is run as a turbidostat without nutrient limitation, the environment selects for both biofilm formation and rapid growth. As shown in Fig. 1, bioreactor-derived lineages have both a strong propensity to form biofilms and faster planktonic growth rates, consistent with the bioreactor selection conditions. Conversely, flask-transfer populations predominately remain in stationary phase and thus, growth rate was not favored. Note that while the bioreactor isolates appear to have reduced cell densities at later time points, the strong biofilms formed by these lineages reduce the accuracy of plate reader cell density readings due to the cells clumping and settling despite shaking. Together these data show that the adaptive environments established by the different experimental evolution approaches favored distinct phenotypes that could reveal potentially different evolutionary trajectories leading to DAP resistance.

### Flask-adapted HOU503 repeatedly evolved mutations within *yvcRS* that led to an increase in *dltABCD* and *mprF* transcripts, consistent with the repulsion phenotype

Comparison of the 10 flask-transfer isolate WGS to HOU503 revealed the repeated evolution of mutations in *orf_*2375 or *orf_*2376, which are annotated as homologs of the *yvcRS* system associated with bacitracin resistance in *Efc* and was recently found to be involved in *Efc* DAP resistance without a functional LiaFSR system (Table 1 and Supplementary Text) (34). Below, we report on the genomic mutations while plasmid dynamics are discussed in the Supplementary Text. YvcS is a transmembrane permease that senses bacitracin and, in conjunction with YvcR, a cytoplasmic ATPase, transmits the signal to the YxdK sensor kinase and YxdJ response regulator (34). This system shares significant similarity to the VraFG/GraSR system in *S. aureus* that upregulates both *dltABCD* and *mprF* in the presence of cationic antimicrobial peptides (CAMPs) (17). Interestingly, *yvcRS* in enterococci is located directly upstream of the *dltABCD* operon. Therefore, we hypothesized that, in *Efm*, YvcRS could regulate *dltABCD* and possibly *mprF.*

**Table 1:** Flask-transfer isolate genomes.

| **Isolate** | **DAP**<br>**MIC** | **VAN**<br>**Plasmid** | ***orf_2376*** | ***orf_2375*** | ***cls*** | ***gdpD*** | ***pgsA*** | ***orf_1482*** |
| --- | --- | --- | --- | --- | --- | --- | --- | --- |
| **FT1** | **32** | **Δ** | **G133V** |  |  |  | **-62** |  |
| **FT2** | **32** | **Δ** | **G133V** |  |  |  | **-62** |  |
| **FT3** | **32** | **Δ** | **T156I** |  |  |  |  | **F130L** |
| **FT4** | **32** | **Δ** | **T156I** |  |  |  |  | **F130L** |
| **FT5** | **32** | **Δ** | **S23I** |  | **R218Q** |  |  |  |
| **FT6** | **32** |  |  | **H203Y** | **A20D** |  |  |  |
| **FT7** | **17** | **Δ** | **I576N** |  |  | **H29R** |  |  |
| **FT8** | **32** | **Δ** | **I576N** |  |  | **H29R** |  |  |
| **FT9\*** | **32** | **Δ** | **T156I** |  |  |  |  | **F130L** |
| **FT10** | **32** | **Δ** |  | **G173C** | **A20D** |  |  |  |
| **Total Strains**<br>**with Changes** |  | **9** | **7** | **2** | **3** | **2** | **2** | **2** |

\*Additional changes may be present within this genome. See Supplementary Text.

To test this hypothesis, we performed qPCR to measure the effect of *yvcRS* mutations on the transcription of *dltA* and *mprF*. As shown in Fig. 2, flask-transfer isolates had 2-9 fold increased *dltA* transcripts and 2-5 fold *mprF* transcripts when compared to the housekeeping gene: glucose-1-dehydrogenase 4, *gdhIV*. Furthermore, all *dlt* operon transcripts were upregulated in FT6 (*yvcR*^H203Y^, *cls*^A20D^) suggesting a link between the *dlt* operon and YvcRS (Fig. S1).

**Fig. 2:**
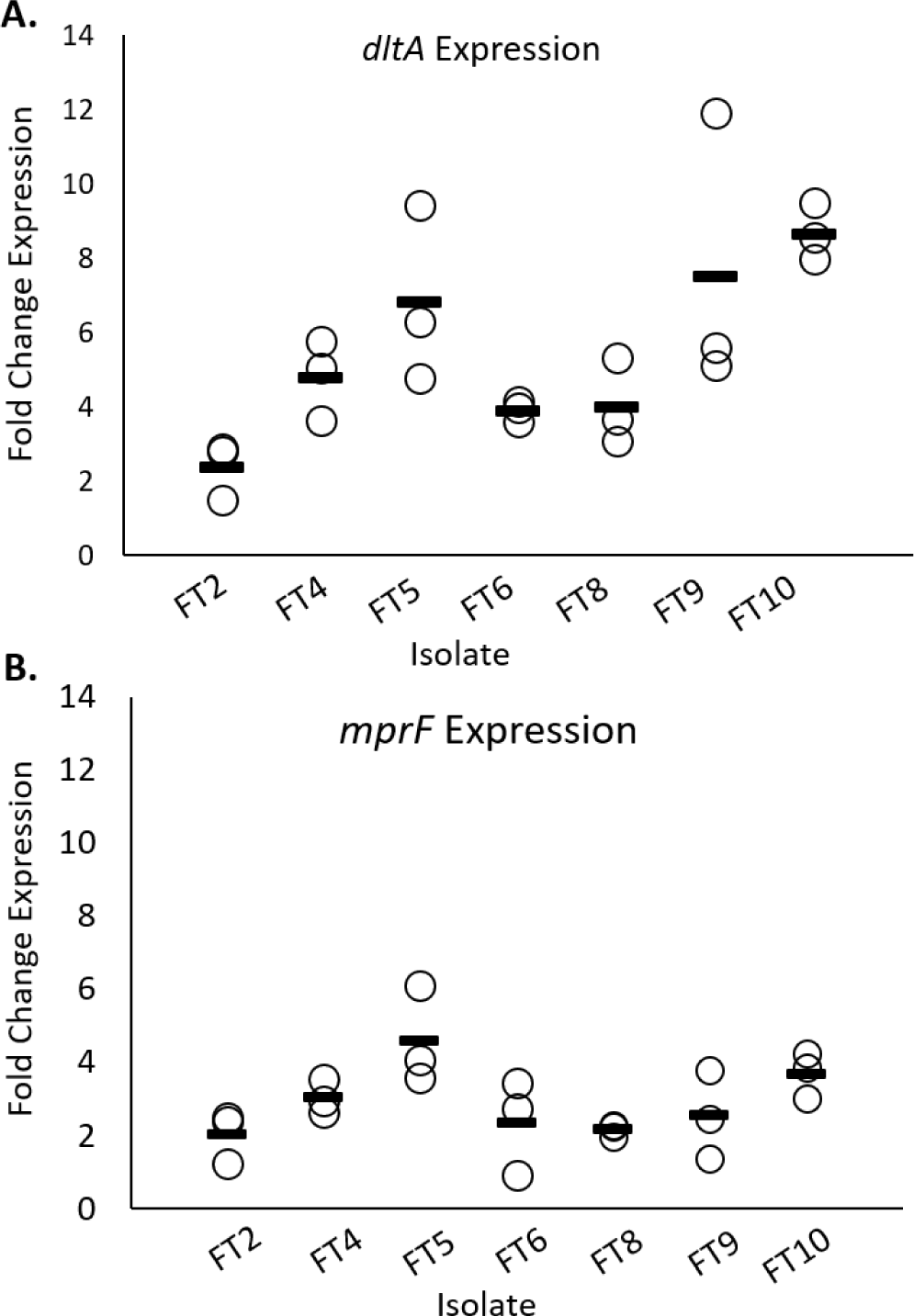
Flask-transfer isolates with mutations in *yvcRS* had upregulated *dltA* and *mprF* transcripts. qPCR was used to measure transcript levels of flask-transfer isolates using *gdhIV* as reference. **A.** *dltA* transcript levels compared to ancestor. **B.** *mprF* transcript levels compared to ancestor.

The isolates used for the assays in Fig. 2 and below contained one additional SNP outside *yvcRS*, making it difficult to assert causality for the mutations in *yvcRS* and the upregulation of the *dltABCD* and *mprF* transcripts. We report the genetic changes associated with DAP resistance within metagenomic analysis of the populations but with the caveat that other mutations can be present within the end-point isolates. For example, to evaluate the likely effects of *yvcS* mutations, we characterized isolates FT2 (*yvcS*^G133V^, *pgsA*^-62^) and FT5 (*yvcS*^S23I^, *cls*^R218Q^) because the only common mutation between them was present in *yvcS*, suggesting that the *yvcS* mutations caused the increase in transcripts. Similarly, FT6 (*yvcR*^H203Y^, *cls*^A20D^) and FT10 (*yvcR*^G173C^, *cls*^A20D^) were chosen to study the effects of *yvcR* mutations as the secondary mutation, *cls*^A20D^, is in the well-characterized *cls* gene. Mutations in *cls* are commonly found in the enterococcal DAP response and are associated with phospholipid “redistribution” (13, 35) and as shown in our later analyses, other lineages containing *cls*^A20D^ had a strikingly different phenotype, suggesting that *cls*^A20D^ could not be directly responsible for the phenotype observed here. By comparing strains with different secondary mutations, the basis of the conclusion linking potential upregulation of *dltABCD* or *mprF* is one of parsimony rather than direct causality.

### Isolates with mutations in *yvcRS* had a more positively charged cell surface and bound less BDP:DAP than the ancestor

To determine if isolates containing *yvcRS* mutations possessed an increase in cell surface charge due to the upregulation of the *dltABCD* operon and *mprF*, flask-transfer isolates were incubated with the positively charged molecule, Poly-L-Lysine, conjugated to FITC (PLL:FITC) and cell fluorescence was quantified using fluorescence microscopy. Cells binding less PLL:FITC correlates with a more positively charged cell surface (36). All flask-transfer isolates tested bound 30-60% less PLL:FITC than the ancestor (p<0.05 using two-sided t-test), indicating that flask-transfer isolates produced a more positively charged cell surface and suggesting that the regulatory changes shown in Fig. 2 may lead to an increase in cell surface charge (Fig. 3A and Fig. S2). We also examined whether the isolates with mutations in *yvcRS*, bound less bodipy-DAP (BDP:DAP) than the ancestor. BDP:DAP is a conjugation of the fluorophore, bodipy-FL, with DAP (BDP:DAP) and is used as a proxy for DAP binding (27, 28, 37). After incubation with 32 µg/ml BDP:DAP, bound BDP:DAP was quantified using fluorescence microscopy (Fig. 3B-C). All flask-transfer isolates bound 53-70% less BDP:DAP than the ancestor (p<0.05 using two-sided t-test), supporting the hypothesis that mutations in *yvcRS* are associated with repulsion of DAP from the cell surface. Incubation with the anionic phospholipid dye, 10-N-nonyl acridine orange (NAO) (15, 38, 39) revealed no redistribution of phospholipid microdomains (Fig. S3).

**Fig. 3:**
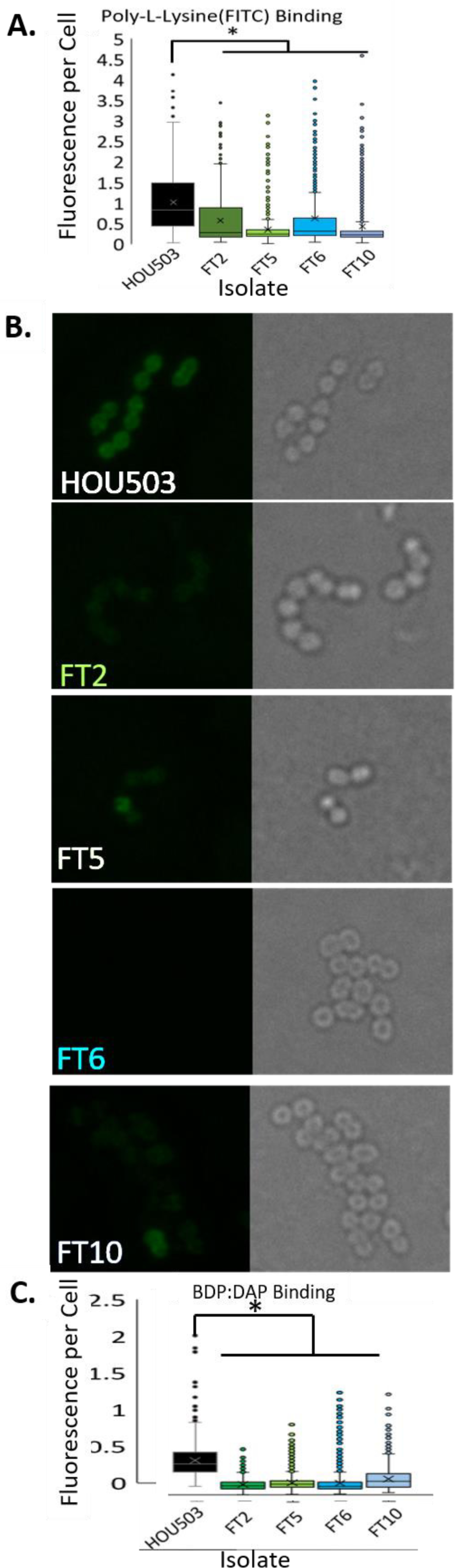
Isolates with mutations in *yvcRS* had a more positively charged cell surface and bound less BDP:DAP than the ancestor, HOU503. **A** The relative cell surface charge was determined by incubation with PLL:FITC. Cells that bind less PLL:FITC have a more positive cell surface charge. *Shows statistical significance (p<0.05) using two-sided T-test. ImageJ was used for quantification. Physical images can be viewed in Fig. S2. **B.** Isolates were incubated with BDP:DAP to determine DAP binding patterns. **C.** Quantification of BDP:DAP binding per cell. *Shows statistical significance (p<0.05) using two-sided T-test. ImageJ was used for quantification.

### Adaptation in a bioreactor produced two main DAP resistance trajectories via mutations in *divIVA* or *oatA*

Using methods described previously (25, 30–33) and more extensively in the Supplementary Text, two independent HOU503 populations were evolved to DAP resistance within 12 and 10 days (Population 1 and Population 2, respectively) in a bioreactor system that favors polymorphic populations and biofilm formation (25, 30, 31, 33). To identify the frequency of each mutation over time and the likely order of mutations, a polymorphic sample was taken daily for metagenomic deep sequencing (Fig. S4). Afterwards, 10 and 9 phenotypically diverse end-point isolates from Population 1 and 2, respectively, underwent WGS to identify the linkages between mutations (Tables 2-3). These end-point isolates were selected based upon phenotypic diversity to increase the chance of identifying diverse genotypes and, therefore, the percentage of end-point isolates carrying any particular mutation does not represent the frequency of that mutation within the entire population. By combining the daily frequencies with the genetic linkages from end-point isolates, adaptive timelines were created, detailing the likely sequence of events that resulted in DAP resistance (Fig. 4). Note that these trajectories do not include plasmid-associated mutations. Plasmids are easily transferred horizontally and, thus, pinpointing a plasmid-encoded mutations’ acquisition in different lineages is difficult (Supplementary Text)(40). Interestingly, several identical mutations were present on Day 1 of both populations (*rpoB*^G32G^, *purA*^L409S^, *ansP*^-288^, *tagB*^R330S^, *orf_*280^C84C^, *orf_*2338^*-*276^), suggesting heterogeneity at these loci in the ancestor. Because these mutations were present at high frequencies on Day 1, without DAP present, and their frequency fell upon the addition of DAP, it is likely that these mutations were not the result of DAP adaptation and were likely hitchhiker mutations. Thus, any mutations identified prior to the addition of DAP were removed from subsequent analysis.

**Table 2:**
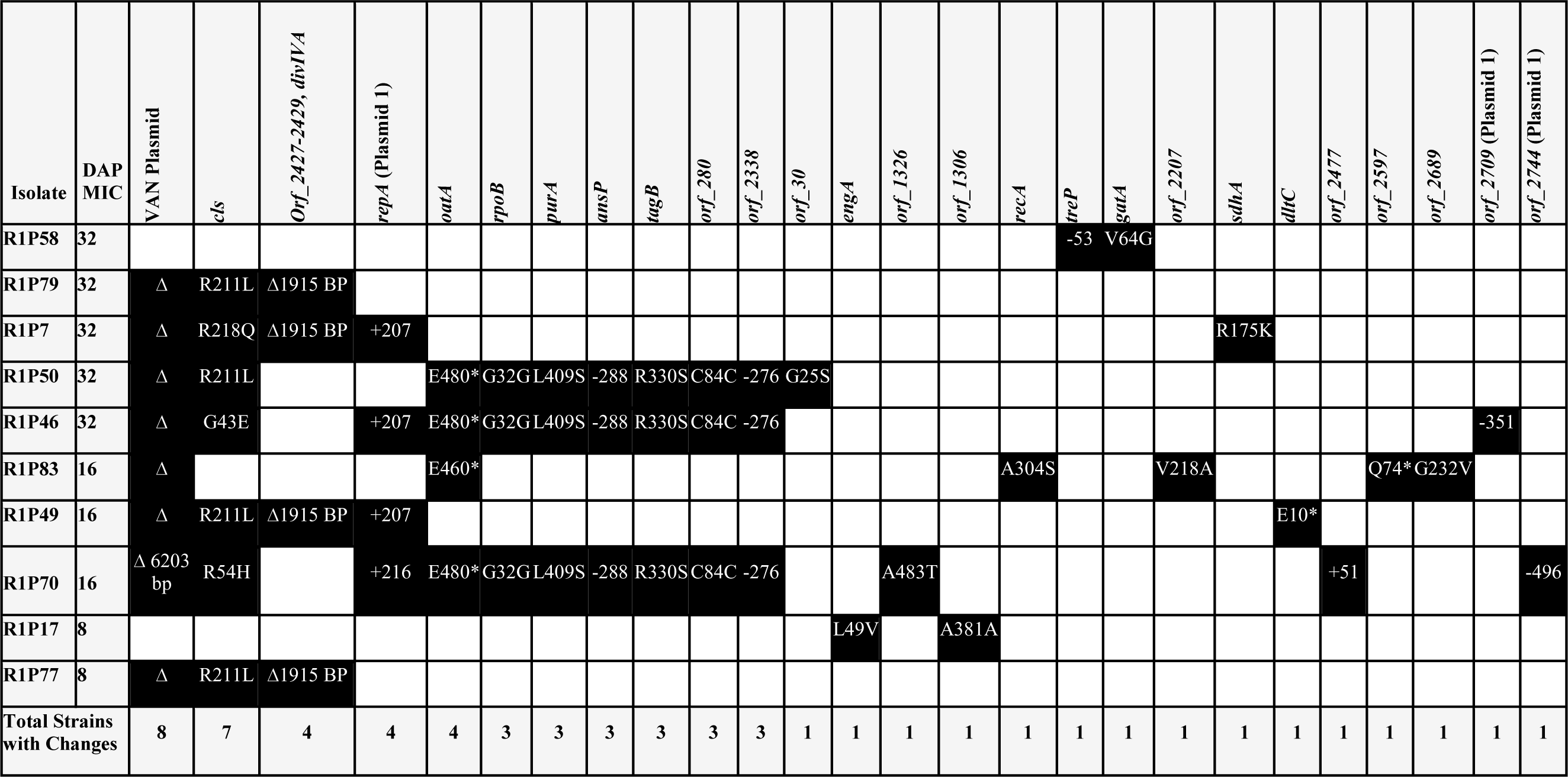
Bioreactor-derived Population 1 end-point isolate genomes.

**Table 3:** Bioreactor-derived Population 2 end-point isolate genomes.

| Isolate | DAP MIC | *divIVA* | *cIs* | *orf_771* | von Willebrand Factor A homologue | *perR* | *Orf_923* | *orf_1323* | *recA* | *rpoB* | *Orf_2201* | *mraY* | *orf_2504* | *orf_2563* | *Orf_2597* | *orf_2902* (Plasmid 1) | *repA* (Plasmid 1) | VAN Plasmid | *orf_3019* (Plasmid 3) |
| --- | --- | --- | --- | --- | --- | --- | --- | --- | --- | --- | --- | --- | --- | --- | --- | --- | --- | --- | --- |
| R2P90 | 64 | Q75K | A20D |  |  |  |  |  |  |  |  |  |  |  |  |  |  |  |  |
| R2P6 | 64 | Q75K | R218Q |  |  |  |  |  |  |  |  |  |  |  |  |  |  |  | K334* |
| R2P74 | 64 | Q75K | A20D |  | D138D |  |  |  |  | -153 |  |  |  |  |  | Q249H |  |  |  |
| R2P76 | 32 | Q75K | A20D |  |  |  |  |  |  |  |  |  |  |  |  |  |  |  |  |
| R2P59 | 32 | Q75K |  |  |  |  | Δ18 bp | E222K |  |  |  |  |  | F227* |  |  |  |  |  |
| R2P61 | 32 | Q75K | R211L |  | E342K |  |  |  |  |  |  | G214D | -5 |  |  |  |  |  |  |
| R2P29 | 16 |  |  | -626 |  |  |  |  |  |  |  |  |  |  |  |  |  |  |  |
| R2P63 | 16 | Q75K | H215R |  | E342K |  |  |  |  |  |  |  |  |  |  |  | +207 | Δ |  |
| R2P91 | 8 |  |  | -626 |  | G109E |  |  |  |  |  |  |  |  |  |  |  |  |  |
| Total Strains with Changes |  | 7 | 6 | 2 | 3 | 1 | 1 | 1 | 1 | 1 | 1 | 1 | 1 | 1 | 1 | 1 | 1 | 1 | 1 |

**Fig. 4:**
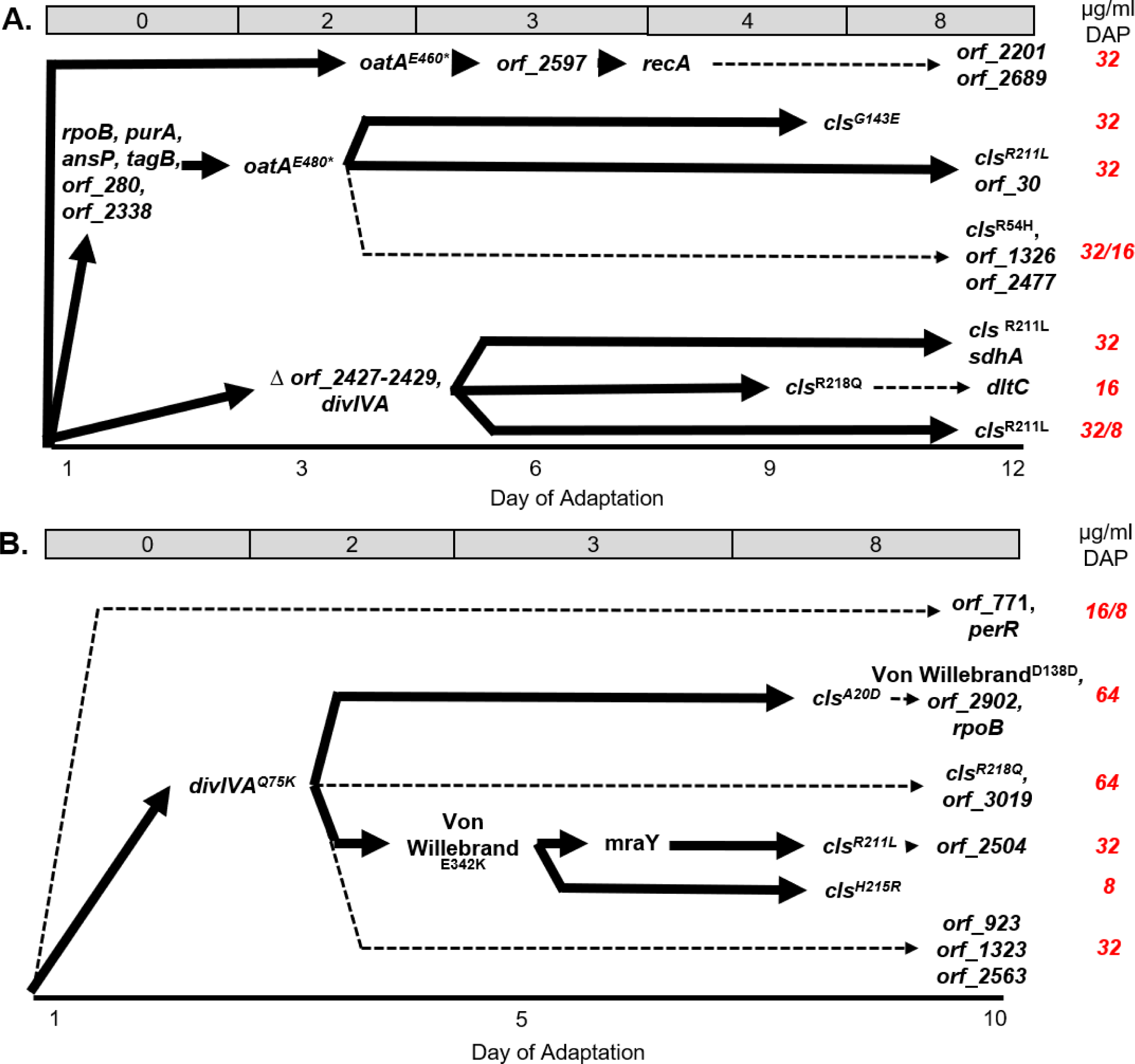
Adaption within a bioreactor environment favoring rapid growth and biofilm formation produced two predominant evolutionary trajectories. Combining the WGS data from end-point isolates that identified genetic linkage with the metagenomic frequency data over time, established the likely sequence of events that resulted in DAP resistant trajectories. Dashed lines indicate that the frequency of the subsequent mutation(s) identified in specific end-point isolates were below the level of detection (<3%) in the overall bioreactor population. The low frequency of these mutations within the population suggests that they were acquired towards the end of adaptation. The DAP concentration each day of the experiment is across the top, in gray. Final DAP MICs of each trajectory are denoted in red at the right. **A.** Population 1 evolved 2 main trajectories opening with either a mutation in *oatA* or Δ1915 followed by additional mutations, including *cls.* **B.** Population 2 evolved one main trajectory with a mutation in *divIVA* followed by mutations in *cls.*

Because mutations that arise early and at high frequency are those more likely to contribute to DAP resistance (25, 31–33), we identified *divIVA* and *oatA* as candidate genes for further study (Fig. 4). A common evolutionary trajectory within Population 1 was Δ*orf_*2427-2429, *divIVA*: a deletion of 1915 bps (hereafter referred to as Δ1915) that resulted in the truncation of the C-terminus of a putative *sepF* gene (residues 107-200), deletions of an S4 domain containing gene and a *yygT* superfamily gene, and the N-terminus deletion of *divIVA* (residues 1-75). SepF forms ring structures at the division septum and directly interacts with FtsZ in *B. subtilis* (41, 42), while DivIVA acts as a scaffold in the septum formation complex and aids in chromosomal segregation by providing a scaffold at the cellular poles (43). In Population 2, a SNP in *divIVA* (Q75K) was the prominent allele, found in 69% of the population, suggesting the importance of changes in *divIVA* towards DAP resistance. Alternatively, mutations in *oatA* were observed in four end-point isolates in Population 1 and were prominent alleles, based on daily frequency data, during early adaptation in Population 2.

After the acquisition of the primary mutations above, mutations in *cls* were acquired and found in 13/19 end-point isolates comprising 36% and 61% of the final populations. The emergence of mutations in *cls* closely mimics *Efc* adaptation to DAP, corroborating the important role of Cls in the evolution of DAP resistance, but only after an initial set of mutations establish a biochemical basis for their acquisition (25). See Supplementary Text for further allelic discussion.

### Bioreactor-derived isolates containing *divIVA* associated mutations produced abnormal division septa

In total, 11 out of 19 bioreactor-derived isolates contained a mutation in *divIVA* that comprised 41% and 69% of the final day populations, respectively (Fig. 4, Tables 1-2, Fig. S4). In Population 1, Δ1915 affected four genes, all of which had predicted functions involved in cell division, including N-terminal deletion of *divIVA*. The mutation observed in Population 2 (Q75K) was in a predicted loop region of *divIVA* between the first two predicted helices of the N-terminal domain. Because the deletion affecting *divIVA* would result in a loss of function, we speculate that *divIVA*^Q75K^ had reduced function, though this remains untested. Interestingly, *divIVA*^Q75K^ was also identified in Population 1, though it was less successful than Δ1915 (Fig. S4A). To understand how Δ1915 affects DAP resistance, we evaluated isolate R1P79 (Δ1915, *cls*^R211L^) as the only additional genomic mutation was *cls*^R211L^, which was also observed in a separate trajectory not containing mutations within *divIVA*: R1P50. To study *divIVA*^Q75K^, we used isolate R2P90, containing *divIVA*^Q75K^ and *cls*^A20D^ (the *cls* allele assessed in FT6 and FT10).

Using transmission electron microscopy (TEM), we found that the Δ1915 and *divIVA*^Q75K^ containing isolates had increased abnormal septation events and abnormal clustering or chaining (Fig. 5). We measured the number of cells with abnormal septation events compared to the ancestor to provide a quantitative metric and found that R1P79 (Δ1915, *cls*^R211L^) exhibited an increase in abnormal septation events from 9% (ancestor) to 82%, whereas R2P90 (*divIVA*^Q75K^, *cls*^A20D^) produced 50% abnormal septation events (Fig. 5D). This suggests that Δ1915 produced a more severe phenotype – consistent with losing two additional genes associated with cell division and the truncation of the cell division allele *sepF* in addition to the loss of *divIVA*. It is important to note that isolates containing *cls*^A20D^ and *cls*^R211L^ did not have obvious septal defects in the presence of *yvcRS* mutations (Fig. 3), supporting the observation that changes in *divIVA* are associated with septal abnormalities.

**Fig. 5:**
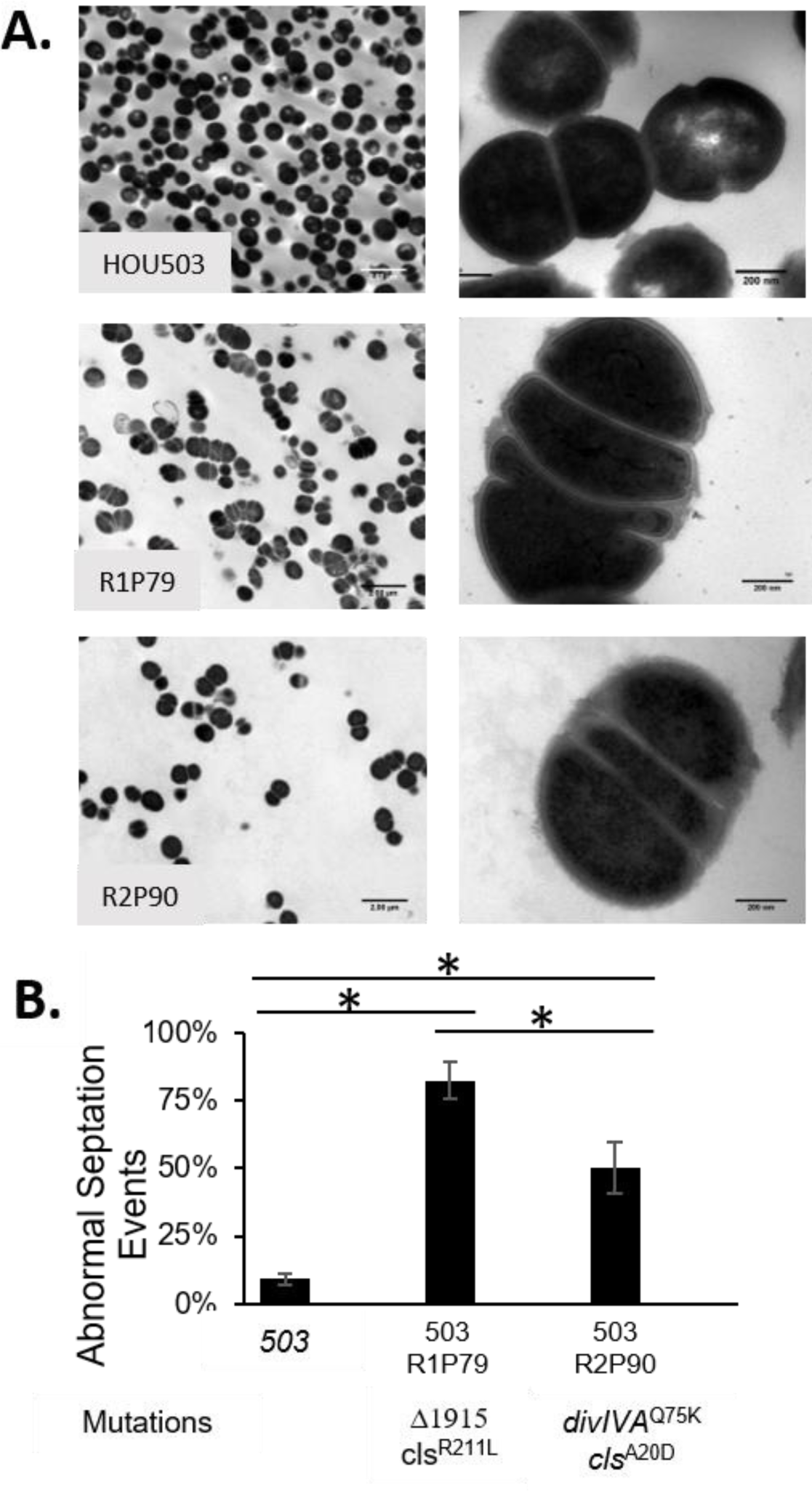
Bioreactor isolates containing *divIVA* associated mutations produced abnormal septa. **A.** TEM was used at 5000x (left) and 50000x (right) to observe cellular morphology of end-point isolates containing mutations in *divIVA.* **B.** The percent of abnormal septal events was determined. *Shows statistical significance (p<0.05) using two-sided T-test.

### Bioreactor-derived isolates containing *divIVA* associated mutations produced differing DAP resistance mechanisms

We assayed PLL:FITC, BDP:DAP, and NAO binding to determine how mutations in *divIVA* resulted in DAP resistance. We found that R1P79 (Δ1915, *cls*^R211L^) bound similar PLL:FITC to the ancestor, yet bound BDP:DAP in a speckled manner, similar to that observed in DAP-resistant *Efc*, R712(44). Incubation with NAO also revealed a speckled phenotype, suggesting that Δ1915 resulted in a redistribution of lipid microdomains and DAP binding (Fig. 6 and Fig. S2). Interestingly, R2P90 (*divIVA*^Q75K^, *cls*^A20D^) while producing aberrant septa showed different staining patterns: binding less PLL:FITC (suggesting a more positive surface), showing no NAO redistribution, and binding BDP:DAP in a strikingly bi-modal manner, reminiscent of streptococci (Fig. 6) (21). Only a subpopulation (approximately 10%) of cells bound significantly more BDP:DAP than the ancestor, whereas the remaining population did not have any discernable difference in drug binding (Fig. 6C). Thus, mechanistically, in R2P90 (*divIVA*^Q75K^, *cls*^A20D^), DAP resistance appears to have been achieved by a combination of cell envelope changes associated with altered cell division and modest changes in cell surface charge resulting in hyperaccumulation of DAP in a subset of cells.

**Fig. 6:**
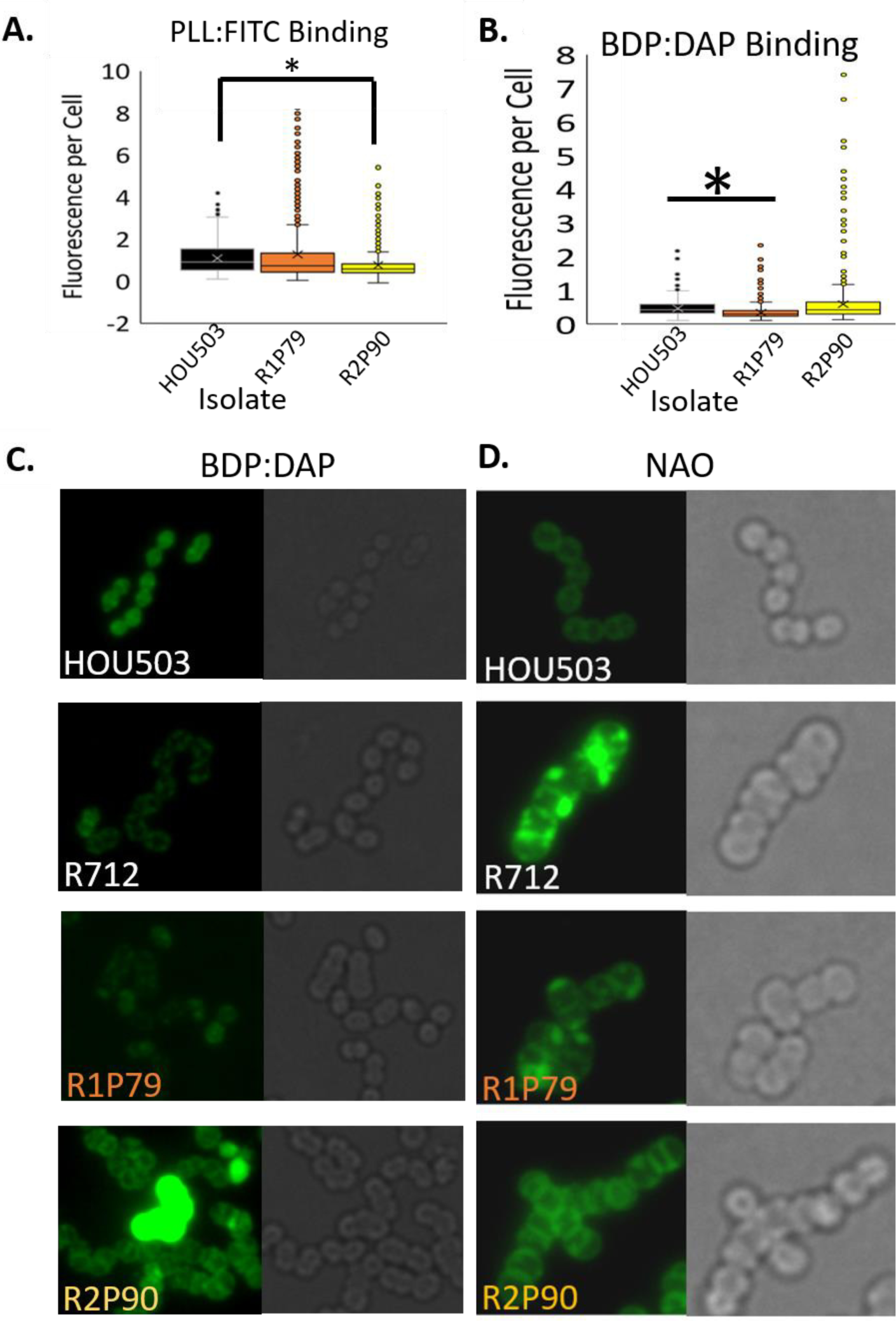
Bioreactor Isolates with *divIVA* associated mutations produced more complex DAP resistance phenotypes. **A.** The relative cell surface charge was determined by incubating the isolates with PLL:FITC Cells that bind less PLL:FITC have a more positive cell surface charge R2P90 bound significantly less PLL:FITC than the ancestor indicating a more positively charged cell surface. *Shows statistical significance (p<0.05) using two-sided T-test. Physical images can be viewed in Supplementary Fig. 2. **B.** Quantification of BDP:DAP binding per cell using ImageJ. *Shows statistical significance (p<0.05) using two-sided T-test. **C.** Isolates were incubated with BDP:DAP to determine DAP binding patterns. *EEfc* R712 acts as a control to show the redistribution of binding phenotype. **D.** Isolates were incubated with NAO to determine phospholipid microdomain patterning. *Efc* R712 acts as a control to show the redistribution phenotype.

### Bioreactor-derived isolates with mutations in *oatA* had decreased peptidoglycan O-acetylation and increased cell surface charge, consistent with reduced BDP:DAP binding

Seven different *oatA* mutations were observed between Bioreactor Population 1 and Population 2, six of which resulted in a truncation of the catalytic C-terminal domain (predicted residues 460-628). OatA catalyzes the acetylation of the C-6 hydroxyl group of *N*-acetylmuramic acid (MurNAc), which contributes to lysozyme resistance and is linked to increased pathogenesis in many species (45–47). Here, two different mutations were identified in end-point isolates: E480* and E460* (Table 2). All *oatA* mutations, combined, comprised 29% of the Population 1 final day (Fig. S4). While *oatA* mutations were not observed in Population 2 end-point isolates, two separate *oatA* mutations (E598* and E460*) were prominent throughout the Population 2 experiment (Fig. S4). Their combined presence remained at over 50% of the population between Days 5-7, but fell to 8% of the final day, as the mutation, *divIVA*^Q75K^, found greater success.

The isolates containing mutations in *oatA* had several additional mutations. We selected R1P50 (*oatA*^E480*^, *cls*^R211L^, *rpoB*^G32G^, *purA*^L409S^, *ansP*^-288^, *tagB*^R330S^, *orf_*280^C84C^, *orf_*2338^*-*276^, *orf_30*^G25S^) and R1P83 (*oatA*^E460*^, *recA*^A304S^, *orf_2207*^V218A^, *orf_2597*^Q74*^, *orf_2689*^G232V^) for further analysis as the only shared mutation between these two end-point isolates were truncations of *oatA.* Isolates with either *oatA* mutation had increased sensitivity to lysozyme as tested via disc diffusion, suggesting a loss in catalytic activity and a likely decrease in peptidoglycan O-acetylation (Fig. 7A). Note that the high concentration of lysozyme in the two left discs caused the opaque halo around the disc, surrounded by the clearance of cells beyond. Additionally, we found that R1P50 and R1P83 bound 10% and 40% less PLL:FITC on average, respectively (Fig. 7B and Fig. S2) and bound less BDP:DAP (39% and 33%, respectively) than the ancestor (Fig. 7C-D). This suggests that the trajectories with *oatA* mutations conferred increased DAP resistance via repulsion of DAP from the cell surface. Incubation with NAO revealed no evidence of phospholipid redistribution (Fig. S3). Thus, in both planktonic and biofilm-heavy environments, *Efm* was able to repulse DAP.

**Fig. 7:**
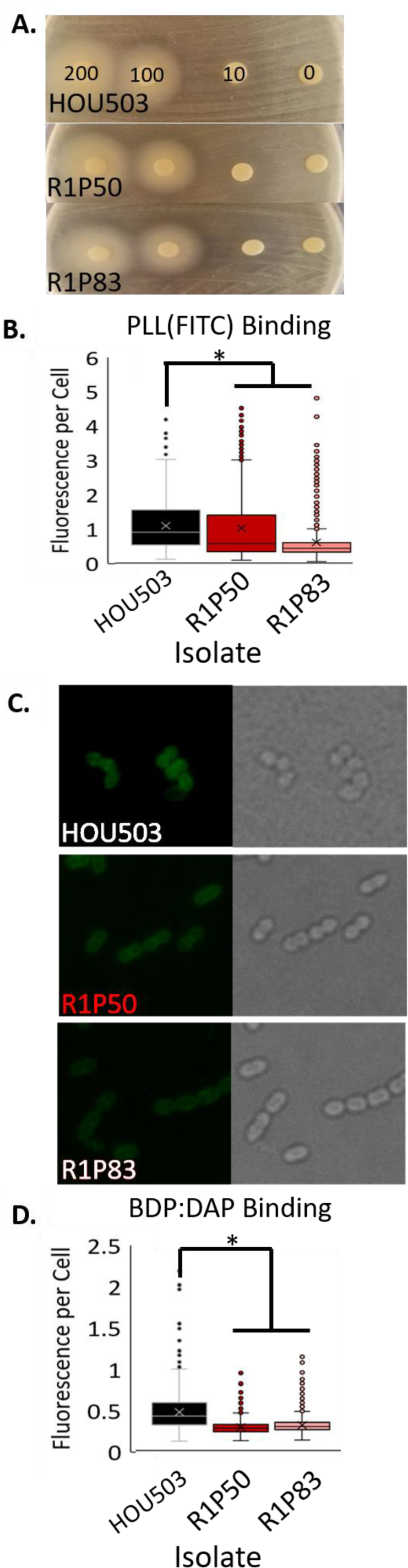
Bioreactor-derived isolates with mutations in *oatA* had decreased peptidoglycan O-acetylation and increased cell surface charge, consistent with reduced BDP:DAP binding. **A.** Lysozyme discs containing decreasing concentrations (200, 100, 10, 0 mg/ml) were overlaid on bacterial lawns. R1P50 and R1P83 both have larger zones of inhibition around lysosome soaked discs indicating a loss in O-acetylation. **B.** The relative cell surface charge was determined by incubating the isolates with PLL:FITC. Cells that bind less PLL:FITC have a more positive cell surface charge. *Shows statistical significance (p<0.05) using two-sided T-test. ImageJ was used for quantification. Physical images can be viewed in Supplementary Fig. 2**. C.** Isolates were incubated with BDP:DAP to determine DAP binding patterns. **D.** Quantification of BDP:DAP binding per cell. *Shows statistical significance (p<0.05) using two-sided T-test. ImageJ was used for quantification.

### After initial changes to establish either repulsion or redistribution, all evolutionary trajectories converged on alleles linked to membrane homeostasis

Regardless of adaptative technique or DAP resistance mechanism, after initial mutations were acquired, changes were made to genes affecting lipid/membrane homeostasis. The most commonly affected gene was *cls*, affecting three flask-transfer isolates, 13 bioreactor-derived isolates, and comprising 36% and 61% of the Population 1 and 2 final days, respectively. In flask-transfer isolates without a mutation in *cls*, mutations in glycerophosphoryl diester phosphodiesterase (*gdpD*), phosphatidylglycerol synthase *(pgsA*), or *orf_1482* (upstream of a different putative *pgsA*, denoted *orf_*1481) were observed, suggesting that while mutations to *cls* were more common, there were alternative evolutionary trajectories that affect membrane phospholipids that could potentially produce the same biological outcome. GdpD is a component of cell membrane phospholipid metabolism and *gdpD* mutations have been reported in clinical DAP resistant isolates *Efm* R494 and *Efc* R712 (I283P and ΔI170, respectively) as well as in one flask-adapted populations here (*gdpD*^*H29R*^) (3, 39). A single G→A mutation was observed 62 base pairs upstream of *pgsA* (*pgsA*^-62^) in FT1 and FT2. PgsA catalyzes the formation of phosphatidylglycerol-3-P from CDP-diacylglycerol and then converted to phosphatidylglycerol by PgpABC. DAP adaptive mutations in *pgsA* have previously been reported in streptococci, *S. aureus*, and *B. subtilis* that result in a loss of catalytic function or decrease in expression, causing a potential decrease in available phosphatidylglycerol for DAP binding (48). Here, we found that isolates containing mutations upstream of *pgsA* and the putative *pgsA* (mutation in *orf_1482* affecting *orf_*1481 expression) produced a very modest decrease in both transcripts, respectively (Fig. S5). It is possible that these transcript reductions result in less phosphatidylglycerol and contribute to DAP resistance.

## Discussion

As MDR bacteria spread, physicians are increasingly forced to administer drugs-of-last-resort, such as DAP, causing resistance to these antibiotics to increase as well. It is predicted that the ascent of pan-resistant strains will result in a ‘post-antibiotic’ era in which many aspects of modern medicine would be threatened. In this work, we have mapped out the DAP resistance trajectories available to *Efm* poised to exploit *liaFSR-*mediated resistance (the most common pathway observed in clinical practice), showing how the environment impacts their acquisition and revealing the multi-layered nature of the resistance phenotype in this organism. Our results provide clarity to the complex and seemingly contradictory sets of observations surrounding the acquisition of DAP resistance in *Efm* and allow us to make important distinctions from *Efc* (5, 27, 39).

Previously, it was found that *Efc* evolves DAP resistance through mutations in the *liaFSR* envelope-stress-response system and *cls* that divert DAP binding away from the division septum of cells via lipid remodeling. Mutations in these systems are seen in the clinic and in *in vitro* studies using both flask-transfers and bioreactors, suggesting that *Efc* evolves DAP resistance largely through phospholipid redistribution (25, 28, 44, 49, 50). Mutations within *Efm liaFSR* have also been observed but strains have not shown evidence of phospholipid redistribution(27). For example, *Efm* HOU503, used here, contains two alleles linked to increased DAP resistance (*liaR*^W73C^ and *liaS*^T120A^) and exhibits tolerance to DAP, but, as shown in Fig. 3B, 6C, 7C, and Fig. S3, neither BDP:DAP nor NAO were redistributed away from the division septa. While *liaR*^73C^ and *liaS*^120A^ may provide HOU503 with an ability to divert DAP, additional mutations were required for HOU503 to survive at higher DAP concentrations. Perhaps surprisingly, none of these additional mutations were present in *yycFG*, a highly conserved two-component system in which mutations have been observed in clinically derived DAP resistant isolates of *Efm* and *S. aureus* (26, 39). This suggests that adapting cells can increase resistance via *liaFSR* or *yycFG* mutations, but not both, implying that the systems could be redundant or engage in cross-talk that produces the same net outputs in signaling. The mutations reported here, after committing to *liaFSR*-associated resistance, predominately resulted in repulsion of DAP from the cell surface. However, in biofilm-heavy and rapid growth environments, HOU503 also diverted DAP binding from septal areas. This combination of resistance strategies marks a major difference between these two closely related species.

In support of the hypothesis that repulsion is a critical driver for DAP resistance in *Efm*, flask-adapted HOU503 repeatedly evolved mutations in the *yvcRS* multi-component system that resulted in the upregulation of the *dltABCD* operon and *mprF*, an increase in cell surface charge, and a reduction in DAP binding. These data suggest that YvcRS in enterococci may be analogous to the VraFG system in *S. aureus* which senses bacitracin and CAMPs in conjunction with GraXSR (potentially analogous to YxdJK in enterococci) to upregulate *dltABCD* and *mprF* and mediate the repulsion phenotype (17, 34, 51, 52).

Notably, our recent study that evolved *Efc* lacking the *liaR* response regulator to DAP resistance in flasks to identify *liaFSR*-independent DAP resistance mechanisms found that the two end-point isolates derived from that experiment contained a mutation in either *yvcR* or *yxdK* (53). These *Efc* isolates containing mutations in the *yxdJK-yvcRS* system did not exhibit an increase in cell surface charge, did not have a reduction in DAP binding, and did not redistribute DAP binding. This phenomenon marks a potential difference between how *yvcRS* functions in enterococcal species, but more importantly, it highlights that when the LiaFSR pathway is disabled, alternate evolutionary trajectories can emerge in *Efc* as is seen here with the multitude of DAP resistance strategies employed by *Efm.* This adaptability also suggests that diverse signaling pathways that respond to environmental stress may be able to act indirectly to compensate for the loss or damage to other systems. Radeck and co-workers noted a similar layered compensatory network in the *B. subtilis* response to bacitracin (54).

In addition to *yvcRS*-mediated repulsion, we also found that changes in MurNAc acetylation through *oatA* mutations contributed to DAP resistance. It is important to note that loss of OatA function, may be illustrative of how repulsion can be achieved, but may be much less likely to occur in a clinical setting. The loss of acetylation leads to lysozyme sensitivity and would likely dramatically decrease pathogenicity, as the host innate immune system would be more effective at clearing this infection (46, 55).

While repulsion was the favored DAP resistance mechanism, redistribution of DAP binding was still observed in a biofilm-heavy environment through the 1915 bp deletion of cell division associated genes, including *divIVA*. The isolate containing this mutation had a dramatic increase in abnormal septation events, redistributed phospholipid microdomains, and diverted DAP binding. The frequent formation of division septa could have the net effect of decreasing the efficiency of DAP in disrupting division by simply increasing the number of potential targets for the antibiotic, ultimately diluting the DAP concentration within the membrane. Interestingly, abnormal septal defects are a common phenotype in DAP resistant isolates of both *Efc* and *Efm* (3, 53).

The role of the mutation *divIVA*^Q75K^ in DAP resistance appears more complex and less clear than the other trajectories. While these cells have a higher frequency of aberrant septa compared to the ancestor, there was no diversion phenotype. Of note, *divIVA*^Q75K^ was the only genotype that produced a strongly bimodal DAP binding phenotype. BDP:DAP stained cells showed a very distinct sub-population that “hyperaccumulated” DAP binding in a uniform manner similar to what has been seen in *Streptococcus mitis/oralis* where it was suggested that such a subpopulation may act as “super-binders” to bind significantly more DAP and thereby protect the surrounding population (21).

After acquiring the mutations that led to either repulsion or DAP diversion, mutations were subsequently acquired in *cls* or other genes associated with lipid/membrane chemistry. *Cls* mutations have been reported in DAP resistant isolates of both *Efm* and *Efc* (3, 25, 27). Here, we found *cls* mutations in both adaptive environments, showing their importance in contributing to DAP resistance. While a variety of mutations in *cls* were observed in the bioreactor (six), two of these mutations were also found in flask-adapted end-point isolates. Both H215R and R218Q have been found in clinical DAP resistant isolates of *Efm* (3). Previous work found that these *cls* mutations increase catalytic activity, creating more cardiolipin, but are neither necessary nor sufficient to confer DAP resistance (35, 39). Furthermore, we found that *cls* mutations were acquired after initial mutations in the *liaFSR* operon during *Efc* DAP adaptation (25). During *Efm* adaptation to DAP in the bioreactor, we again observed that changes to *cls* were acquired at a later stage, suggesting an ordered pathway, regardless of whether repulsion or diversion were being employed as the initial steps (25). While not directly conferring DAP resistance, it is clear that *cls* mutations play an important role in the enterococcal counterattack against DAP. The prevalence of these mutations in the clinic, across species (56), across adaptive environment, and across DAP resistance mechanisms suggests that these alleles may act as good diagnostic DAP resistance markers in clinical infections.

In summary, we have shown here that the environment influences how *Efm* evolves DAP resistance, even with the presence of alleles that are associated with DAP diversion. More planktonic environments select for repulsion-mediated resistance via mutations to the *yvcRS* system that seem to play a role in *dltABCD* and *mprF* regulation. Conversely, environments that favor biofilm and more complex structured communities produce both repulsion and DAP diversion mechanisms, though DAP diversion appears less frequently. It is possible that these selective environments are representative of distinct *Efm* infection environments and may predict how different infections (i.e. bacteremia v. colonization of catheters/stents) will evolve DAP resistance. The unifying theme across resistance strategies, adaptive environments, and enterococcal species was mutations affecting membrane architecture, specifically in *cls*, which may have utility as a DAP resistance marker for clinicians.

## Materials and Methods

### Flask-transfer adaptation

Five populations of HOU503 were adapted to DAP resistance using 100-fold dilutions each day. To start, five different colonies were used to inoculate each of the five populations containing Brain Heart Infusion (BHI) and Ca+ (50 mg/L CaCl_2_). The following day, the populations were transferred to fresh tubes containing 1.5 µg/ml DAP (half the initial DAP MIC). After this, the populations were transferred to two tubes, containing either 1.5× or 2× the current DAP concentration. The tube with the best growth was then propagated into two new tubes. This model was followed until the populations were growing in 8 µg/ml DAP. At the end of adaptation, each population was serially diluted onto non-selective BHI. Two isolates from each population were selected at random for WGS and further analysis.

### Directed evolution of *E. faecium* in a bioreactor

Clinical isolate, *E. faecium* HOU503, was adapted to DAP resistance in two replicate runs in BHI with supplemented calcium (50 mg/L CaCl_2_). Experiments were completed in a Sartorius Stedum Biostat B Plus 1L vessel. A 200 ml culture volume was maintained which received an airflow of 0.2 lpm and was stirred at 100 rpm. The bioreactor was run as a turbidostat, maintaining constant cell density. However, due to the prevalence of biofilm in enterococcal cultures, optical density (OD) probes were rendered useless. To circumvent this problem, CO_2_ was measured by a Magellan Tandem Pro Gas Analyzer and used as a proxy to monitor cell density and maintain logarithmic growth as described previously (25, 30, 31, 33). Manual OD measurements were taken periodically to ensure appropriate cell density. For inoculation, an overnight culture (ON) was grown from a single colony on non-selective media. 1 ml of this ON was then used for inoculating the vessel. The culture was initially grown in the absence of DAP to allow the cells to acclimate to the vessel. DAP was then added at half the initial MIC (1.5 µg/ml). Every two days, MIC testing via two-fold broth dilution was performed on a sample taken from the bioreactor to determine the subsequent DAP concentration. The DAP concentration was only increased in the vessel if the sample culture grew equally well in the higher DAP concentration as was observed in the current, working DAP concentration. By maintaining the population at sub-inhibitory levels of DAP, there is less selective pressure acting on the population, preventing a bottle-neck and allowing for more polymorphism within the population. Samples were taken daily and plated onto BHI and Bile Esculin Agar (BEA) to ensure that the vessel was not contaminated. 3-15 ml samples were taken daily and stored as pellets in −80°C alongside their corresponding supernatants and a glycerol stock. At the end of each run, the population within the vessel was serially diluted and plated onto non-selective BHI agar. To identify phenotypic differences embodying potential different genetic trajectories, 90 isolates were chosen at random and underwent three phenotypic screens: 1) DAP MICs were determined via broth dilution, 2) cell densities at stationary phase were measured, 3) and propensity to grow as floc in broth was noted. Based on these three characteristics, 10 or 9 isolates from each run with diverse characteristics were selected for further characterization and WGS. For further details, see the Supplementary Text.

### Crystal violet biofilm assay

ONs were used to inoculate fresh trypticase soy broth (TSB)and outgrown to OD_600_ 0.5. These cultures were then used to inoculate TSB and grown for 16 hours at 37°C in a 96-well plate with shaking. Planktonic cells were aspirated, and the remaining biofilm was fixed with 99% methanol. Plates were washed three times with PBS and then air dried. The biofilm was stained with 0.2% crystal violet and incubated at room temperature for 15 minutes. The crystal violet was removed followed by three additional PBS washings and allowed to air dry. Bound crystal violet was solubilized in 33% acetic acid and absorbance was measured at *A*_570_. The assay was performed in triplicate.

### Growth rates

ONs were normalized to OD_600_ 0.05 and used to inoculate fresh BHI in a 96 well plate. Cells were grown within a BioTek Epoch2 microplate reader with orbital shaking at 37°C. Measurements were taken every five minutes for 24 hours. Assay was performed in biological triplicates.

### Isolating gDNA and library prep

The UltraClean Microbial DNA Isolation Kit (MoBio) was used for isolating gDNA from both end-point isolates and the daily population samples. Each end-point isolate was grown ON in 10 ml BHI and pelleted. Alternatively, the pellets collected each day from the bioreactor and stored at −80°C were thawed and immediately used for gDNA extraction to eliminate the possible effects of freeze/thaw on the outgrown population. In addition to the published protocol, 5 µL of 5 U/mL mutanolysin and 12.5 µL of 200 mg/mL lysozyme were added to the sample suspended in the Microbead Solution and incubated at 37°C for 1 hour. The Nexterra XT kit was used for the generation of paired-end libraries using 2.5µl gDNA and extending the tagmentation step to 9 minutes at 55°C. Libraries were sequenced by Genewiz on Hiseq with 2×150 bp reads. End-point isolates were sequenced with a minimum 100x coverage and metagenomic sequences were sequenced with at least 300× coverage.

### Analyzing genomic sequencing

Illumina short-reads were aligned to the ancestor, HOU503, using the Breseq-0.29.0 pipeline. Daily samples were analyzed utilizing the polymorphism command (-p) to identify the frequency of each mutation on a given day. Alleles that reached a minimum of 5% on any day were manually examined to ensure accurate mutation calling.

### qPCR

Total RNA was extracted in accordance with the published Qiagen RNeasy Mini protocol with the addition of a 30-minute incubation at 37°C with mutanolysin and lysozyme to help break open the cells. Samples were DNase I treated in accordance with the Invitrogen protocol, using Taq PCR to confirm the removal of contaminating DNA. cDNA was synthesized using Invitrogen SuperScript III in accordance with manufacturer’s instructions. qPCR was performed using Bio Rad iQ Sybr Green in accordance with manufacturer’s instructions on a Bio-Rad CFX Connect Real-Time System. *gdhIV* was used as the housekeeping gene. Changes in expression were calculated using the 2^-ΔΔ*CT*^ method. Experiments were performed in biological and technical triplicate.

### Poly-L-Lysine-FITC assay

Isolates were grown overnight in BHI and then used to inoculate fresh tubes containing BHI. Cells were grown until they reached an OD_600_ 0.5 and then washed 3 times in HEPES (20 mM, pH 7.0). Cells were then resuspended in HEPES to an OD_600_ 0.1 and incubated with 10 µg/ml PLL:FITC, shaking at room temperature for 10 minutes. The culture was then washed once with HEPES to remove unbound PLL:FITC. Cells were resuspended in VectaShield and imaged on a Keyence BZ-Z710. Fluorescence per cell was calculated in ImageJ. Experiments were completed in duplicate on separate days.

### BDP:DAP

The conjugation of the fluorophore, Bodipy-Fl, to DAP was performed as described previously (37). In brief, DAP and BDP were incubated, shaking at room temperature in 0.2 M sodium carbonate buffer, pH 8.5, for one hour followed by extensive dialysis against dH_2_0 at 4°C. BDP:DAP was then incubated with different enterococci isolates with known different BDP:DAP binding patterns at different concentrations to confirm appropriate labeling. To determine BDP:DAP binding patterns, we used methods described previously (11, 12, 44, 57). Briefly, overnights of each isolate were used to inoculate fresh BHI containing Ca^+^ and grown to OD_600_ 0.5. Cells were then incubated with 32 µg/ml BDP:DAP for 20 minutes in the dark at 37°C with shaking. Cells were washed once with HEPES (20 mM, pH 7.0). The labelled pellet was resuspended in VectaShield and mounted onto Poly-L-Lysine coated coverslips and imaged on a Keyence BZ-Z710 using a standard fluorescein isothio-cyanate (FITC) filter. Experiments were completed in duplicate on two separate days. Fluorescence per cell was calculated using ImageJ.

### 10-N-Nonanyl acridine orange (NAO) staining

NAO has been shown to preferentially bind anionic phospholipids in cell membranes and has been used previously to show phenotypes of phospholipid redistribution (15, 38, 44). Isolates were grown to early exponential phase (OD_600_ 0.2) in TSB and then incubated with 500 nM NAO at 37°C with shaking in the dark for 3.5 hours. Cells were then washed three times in 0.9% saline, resuspended in VectaShield, immobilized on Poly-L-Lysine coated coverslips, and visualized on the Keyence BZ-Z710 microscope.

### Transmission Electron Microscopy

Selected isolates were grown ON in BHI. 1 ml of culture was pelleted and washed 3 times in 0.1M Millonig’s phosphate buffer. The pellet was then resuspended in 1 ml glutaraldehyde in Millonig’s phosphate buffer and further processed by the University of Texas Health Science Center Electron Microscopy Core. Imaging was performed on a JEOL JEM 1200 EX Electron Microscope. To quantify abnormal septation, a minimum of 25 “events” were selected in a field of view at 5000x and deemed normal or abnormal in appearance. This was repeated for 6 fields of views, resulting in the characterization of at least 125 events.

### Antibiotic cross-sensitivity

All MICs were determined in triplicate via 2-fold broth dilution. Overnight (ON) cultures were grown at 37°C and shaking at 225 RPM. ONs were normalized to OD_600_ 0.05 and 5 μl was used to inoculate 0.5 ml BHI with different concentrations of antibiotics and grown ON with shaking at 37°C. The lowest concentration with no visible bacterial growth was considered the MIC.

### Lysozyme Sensitivity

ONs were used to inoculate fresh BHI and outgrown to OD_600_ 0.5. These cultures were then used to plate a lawn onto BHI plates. Discs containing increasing lysozyme concentrations were overlain onto the lawn and incubated at 37°C for 48 hours.

### Data Availability

All genomic sequences were submitted under PRJNA522390 (https://www.ncbi.nlm.nih.gov/bioproject/?term=PRJNA522390).

## Acknowledgements

This work was supported by National Institutes of Health, National Institute of Allergy and Infectious Diseases grants R01AI080714 to Y.S., K08 AI135093 to W.R.M., K24-AI121296 and R01-AI134637 to C.A.A., and K08-AI113317 to T.T. Funding agencies did not play a role in experimental design, performance or analysis.

A.G.P., H.M., Y.S., and C.A.A. contributed to experimental design and conceptualization. A.G.P., H.M., and A.J.K. completed experiments. W.R.M. and T.T. aided in analysis and data acquisition. A.G.P., C.A.A. and Y.S. contributed to writing the manuscript.

